# An Antioxidant Enzyme Therapeutic for COVID-19

**DOI:** 10.1101/2020.07.15.205211

**Authors:** Meng Qin, Zheng Cao, Jing Wen, Qingsong Yu, Chaoyong Liu, Fang Wang, Fengmei Yang, Yanyan Li, Gregory Fishbein, Sen Yan, Bin Xu, Yi Hou, Zhenbo Ning, Kaili Nie, Ni Jiang, Zhen Liu, Jun Wu, Yanting Yu, Heng Li, Huiwen Zheng, Jing Li, Weihua Jin, Sheng Pan, Shuai Wang, Jianfeng Chen, Zhihua Gan, Zhanlong He, Yunfeng Lu

**Affiliations:** State Key Laboratory of Organic-inorganic Composites, Beijing Advanced Innovation Center for Soft Matter Science and Engineering, College of Life Science and Technology, Beijing University of Chemical Technology, Beijing, 100029, China; Institute of Medical Biology, Chinese Academy of Medical Sciences, Peking Union Medical College, Yunnan Key Laboratory of Vaccine Research Development on Severe Infectious Disease, Kunming, 650118, China; Department of Chemical and Molecular Engineering, Microbiology, Immunology and Molecular Genetics, and the David Geffen School of Medicine, University of California, Los Angeles, CA 90095, U.S.A; Guangdong-Hongkong-Macau Institute of CNS Regeneration, Jinan University, Guangdong, China; Vivibaba, Inc, 570 Westwood Plaza, University of California, Los Angeles, CA 90095

**Author notes:** These authors contributed equally to this work (Meng Qin, Zheng Cao, Jing Wen). Corresponding author. Email addresses: (J. Chen); (Z. Gan); (Z. He); (Y. Lu).

## Abstract

The COVID-19 pandemic has taken a significant toll on people worldwide, and there are currently no specific antivirus drugs or vaccines. We report herein a therapeutic based on catalase, an antioxidant enzyme that can effectively breakdown hydrogen peroxide and minimize the downstream reactive oxygen species, which are excessively produced resulting from the infection and inflammatory process. Catalase assists to regulate production of cytokines, protect oxidative injury, and repress replication of SARS-CoV-2, as demonstrated in human leukocytes and alveolar epithelial cells, and *rhesus macaques*, without noticeable toxicity. Such a therapeutic can be readily manufactured at low cost as a potential treatment for COVID-19.

---

The severe acute respiratory syndrome coronavirus 2 (SARS-CoV-2) has resulted in over ten million COVID-19 cases globally. Broad-spectrum antiviral drugs (e.g., nucleoside analogues and HIV-protease inhibitors) are being utilized to attenuate the infection. However, current management is supportive, and without specific antivirus drugs or vaccine against COVID-19(*1*). While the pathogenesis of COVID-19 remains elusive, accumulating evidence suggests that a subgroup of patients with severe COVID-19 might have cytokine storm syndrome(*2, 3*). Cytokine storm is a serious immune dysregulation resultant from overproduction of cytokines, which often occurs during virus infection(*4*), organ transplant(*5*), immunotherapy(*6*), and autoimmune diseases(*7*), and may result in death if untreated(*8*). Treatment of hyperinflammation and immunosuppression are highly recommended to address the immediate need to reduce mortality(*2*). Current immunosuppression options include steroids(*9*), intravenous immunoglobulin(*10*), selective cytokine blockade (e.g., anakinra(*11*) or tocilizumab(*12*)), and Janus kinase inhibition(*13*).

In light of the findings that elevated levels of reactive oxygen species (ROS) is strongly correlated with inflammation,(*14*) oxidative injury,(*15*) as well as viral infection and replication(*16–18*), we speculate that regulating the ROS level in COVID-19 patients could be effective for the treatment of hyperinflammation, protection of tissues from oxidative injury, and repression of viral replication. As illustrated in **Scheme 1A**, after infection of SARS-CoV-2, leukocytes are attracted to affected sites releasing cytokines and ROS. An increasing ROS level promotes viral replication, causes oxidative injury, and induces cell apoptosis through DNA damage, lipid peroxidation and protein oxidation, which further exacerbates the immune response. As a result, an increasing number of leukocytes are recruited, further releasing ROS and cytokines, resulting in hyperinflammation and cytokine storm syndrome.

**Scheme 1.**
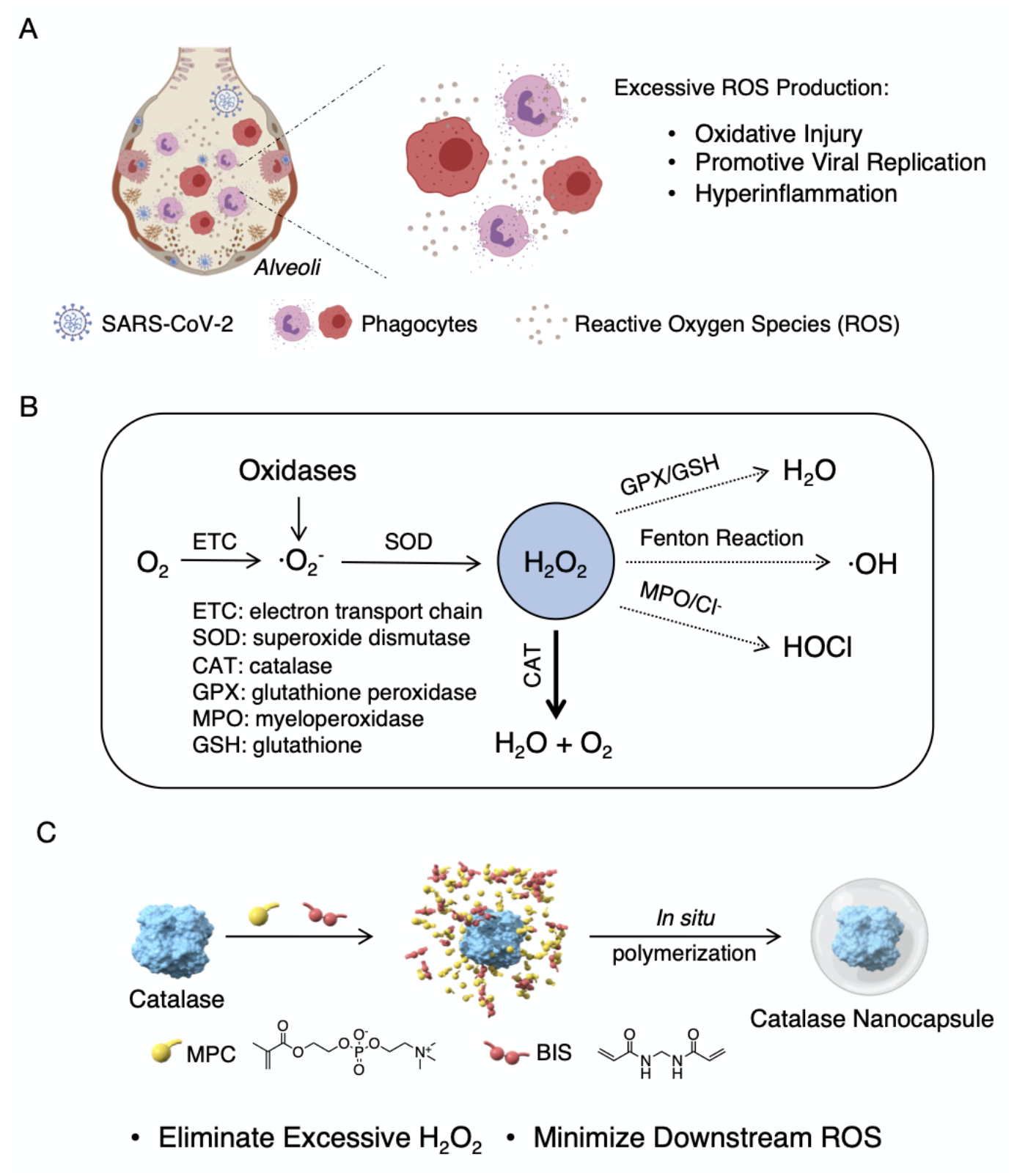
Proposed mechanism of action and synthesis of catalase nanocapsules. (A) A schematic illustrating that an elevated level of ROS causes oxidative injury, promotes viral replication, and triggers cytokine storm syndrome in COVID-19 patients. (B) The reaction pathways of ROS, suggesting that eliminating H2O2 is the key to minimizing the formation of downstream ROS. (C) The synthesis of catalase nanocapsules by *in situ* polymerization of MPC and BIS around individual catalase molecules exhibiting improved stability and circulation halflife.

ROS are a class of partially reduced metabolites of oxygen that possess strong oxidizing capability, which are generated as byproducts of cellular metabolism through the electron transport chains in mitochondria and cytochrome P450(*19*). The other major source are oxidases(*15*) (e.g., NAPDH oxidase), which are ubiquitously present in a variety of cells, particularly phagocytes and endothelial cells. As shown in **Scheme 1B**, partial reduction of O_2_ in these processes generates superoxide anions (·O_2_^-^), which are rapidly converted to hydrogen peroxide (H_2_O_2_) mediated by superoxide dismutase (SOD). H_2_O_2_ may subsequently react forming hydroxyls (OH· and OH^-^) through the Fenton reaction, HOCl through myeloperoxidase (MPO), H_2_O through glutathione/glutathione peroxidase (GSH/GPX), and H_2_O/O_2_ through catalase (CAT), respectively(*19*). Since ·O_2_^-^ possesses a short half-life (~10^-6^ *s*)(*20*), it is rapidly converted to H_2_O_2_, which is chemically stable and able to cross cell membranes and diffuse in tissues. Under a pathological condition, where ROS are excessively produced but antioxidant enzymes are insufficiently presented, H2O2 may accumulate locally or systematically(*21*), which oxidizes proteins with sulfur-containing residues (cysteine and methionine) and reacts with transition metals (e.g., iron), generating downstream ROS that are highly active(*22, 23*). In the context of reaction pathways and kinetics, eliminating the excessive H2O2 is critical to minimize the formation of downstream ROS, prevent oxidative injury, and avoid immunopathogenesis.

Catalase, the most abundant antioxidant enzyme ubiquitously present in the liver, erythrocytes and alveolar epithelial cells, is the most effective catalyst for the decomposition of H_2_O_2_(*24*). One catalase molecule can breakdown 10^7^ H_2_O_2_ molecules in 1 s with an extremely high turnover number of 10^7^ s^-1^; however, catalase generally exhibits poor stability and a short plasma half-life(*25*). To explore its therapeutic use, we encapsulated catalase with a thin shell of polymer through *in situ* polymerization(*26, 27*). As illustrated in **Scheme 1C**, 2-methacryloyloxyethyl phosphorylcholine (MPC), N-(3-aminopropyl) methacrylamide hydrochloride (APM), and N,N’-methylenebisacrylamide (BIS) are used as the monomers and crosslinker. These molecules are enriched around the catalase molecules through noncovalent interactions; subsequent polymerization grows a thin polymeric shell around individual catalase molecules, forming nanocapsules denoted as n(CAT). The thin shell protects the enzyme, while allowing H2O2 to rapidly transport through, endowing n(CAT) with high enzyme activity, augmented stability, and improved plasma half-life.

As shown in **Fig. 1A, B**, n(CAT) shows a size distribution centered at 25 nm and a zeta potential of 1.5 mV, in comparison with those of native catalase (10 nm and – 4.0 mV); TEM image confirms that n(CAT) has an average size of 20~30 nm (**Fig. 1C**). Compared with native catalase, n(CAT) exhibits a similar enzyme activity **(fig. S1A)**, yet with significantly improved enzyme stability. As shown in **Fig. 1E, F**, n(CAT) and native catalase retain 90% and 52% of the activity after incubation in PBS at 37 °C for 24 h, respectively, indicating improved thermal stability. After incubation in PBS with 50 μg/mL trypsin at 37 °C for 2 h, n(CAT) and native catalase retain 87% and 30% of the activity, respectively, suggesting improved protease stability. In addition, n(CAT) in solution retains 100% of the activity after storage at 4 °C and 25 °C for 3 mo. (**fig. S1B**); after freeze drying, n(CAT) retains more than 90% of the activity (**fig. S1C**). Such characteristics are critical for the transport and distribution of n(CAT).

**Fig. 1.**
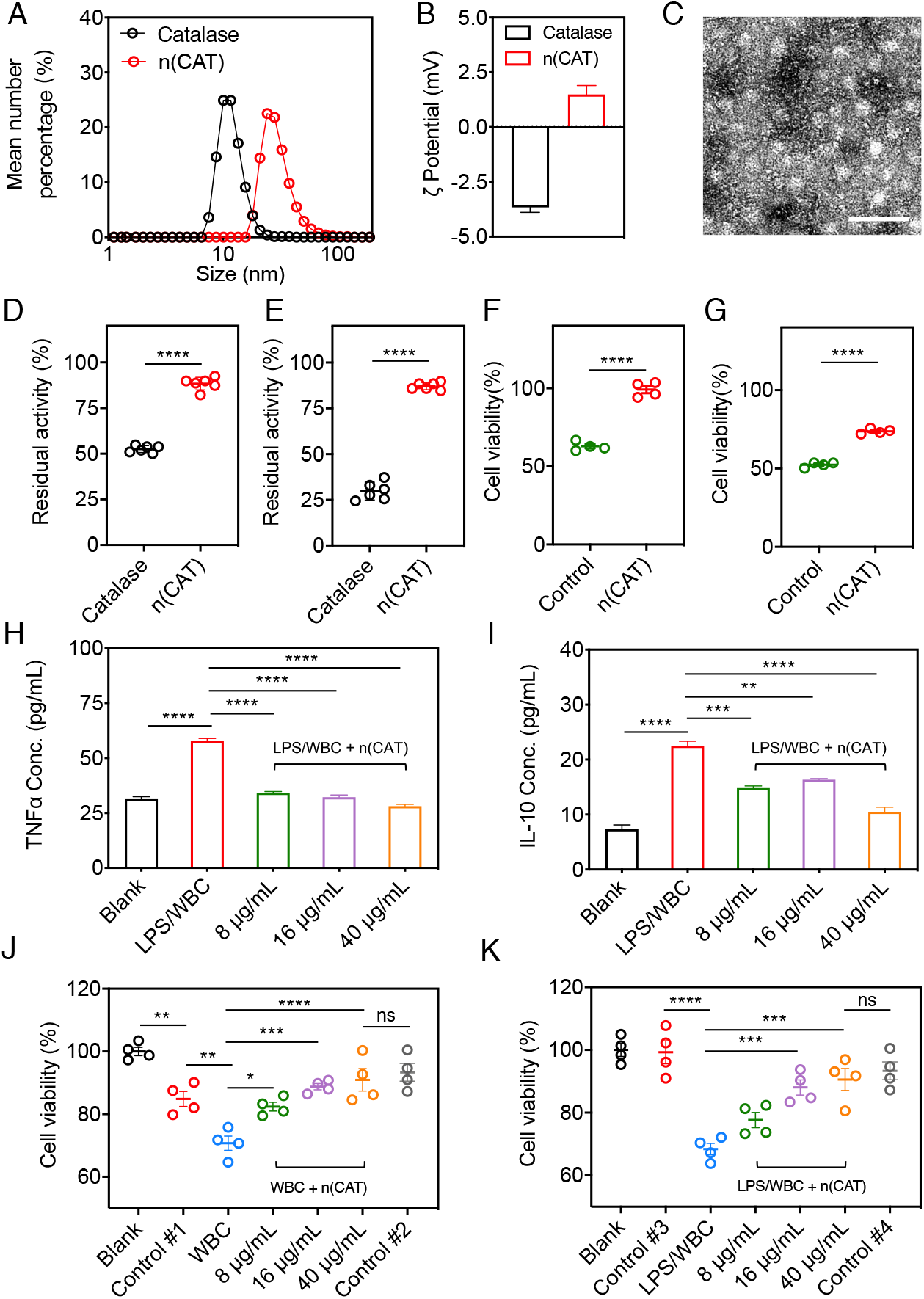
Characteristic, anti-inflammatory effect, and protective ability of n(CAT) (A) Dynamic light scattering; (B) zeta potential; D) thermal stability; (E) proteolytic stability of native catalase and n(CAT). (C) Transmission electron microscopic (TEM) image of n(CAT). (F) Cell viability of HPAEpiC pre-cultured with 20 μg/mL n(CAT) for 12 h, followed by addition of H_2_O_2_ (1000 μM) and culturing for 24 h. (G) Cell viability of HPAEpiC pre-cultured with 1000 μM H_2_O_2_ for 24 h, followed by culturing in fresh media containing 20 μg/ml n(CAT) for 12 h. (H, I) Concentration of (H) TNF-α and (I) IL-10 in the media of human leukocytes (white blood cells, WBC) cultured with LPS and different concentrations of n(CAT). (J) Cell viability of HPAEpiC pre-cultured with 500 μM H_2_O_2_ for 12 h (Control #1) followed by culturing with WBC and different concentrations of n(CAT), as well as that of untreated HPAEpiC cultured with WBC for 12 h (Control #2). (K) Cell viability of HPAEpiC cultured with LPS (Control #3), with WBC (Control 4#), and with LPS, WBC, and different concentrations of n(CAT). *P* value: * < 0.05; ** < 0.01; *** < 0.001; **** < 0.0001.

The ability of n(CAT) to protect lung tissues from oxidative injury was examined in human pulmonary alveolar epithelial cells (HPAEpiC). We first investigated the cytotoxicity of n(CAT) by culturing HPAEpiC with different concentrations of n(CAT) (**fig. S2A)**. The cells with n(CAT) exhibit similar or higher cell viability than the control cells, indicating that n(CAT) does not show any noticeable cytotoxicity to HPAEpiC. The higher cell viability observed is possibly attributable to the ability of n(CAT) to remove H_2_O_2_ produced in the cultures. To examine the protective effect, HPAEpiC were cultured with 20 μg/mL of n(CAT) for 12 h, after which 1,000 μM H_2_O_2_ was added to the media and cultured for 24 h (**Fig. 1F)**. The cells without n(CAT) show a cell viability of 63%, while the cells with n(CAT) retain 100% of the cell viability, demonstrating an ability to protect the cells from oxidative injury. In addition, HPAEpiC were incubated with 1000 μM H_2_O_2_ for 24 h to induce cell injury, after which the injured cells were incubated with 20 μg/mL of n(CAT) for 12 h (**Fig. 1G)**. Culturing the injured cells with n(CAT) increases the cell viability from 50% to 73%, indicating an ability of n(CAT) to resuscitate injured cells. Similar protective and resuscitative effects were also observed with lower n(CAT) concentrations (**fig. S2B, C)**.

Hyperinflammatory response induced by SARS-CoV-2 is a major cause of disease severity and death in patients with COVID-19. The infection and the destruction of lung cells trigger a local immune response, recruiting leukocytes to affected sites(*28*). Unrestrained inflammatory cell infiltration, however, results in excessive secretion of proteases and ROS. In addition to the damage resulting from the virus itself, dysfunctional immune response results in diffusive alveolar damage, including desquamation of alveolar cells, hyaline membrane formation, and pulmonary oedema(*29*). Overproduction of pro-inflammation cytokines is commonly observed in COVID-19 patients, in whom the severity is strongly correlated to the level of cytokines, such as tumor necrosis factor a (TNF-a) and interleukin 10 (IL-10)(*30, 31*). Regulating the production of cytokines, in this context, is critical to reinstate immune homeostasis, and anti-cytokine therapy (e.g., TNF-a antagonist) has been suggested for alleviation of hyperinflammation in severe cases(*32*).

In light of these findings, the ability of n(CAT) to regulate cytokine production was studied in human leukocytes (white blood cells, WBC). Leukocytes were cultured with lipopolysaccharides (LPS, a bacterial endotoxin that activates leukocytes) with and without n(CAT). **Fig. 1H, I** show the concentration of TNFa and IL-10 in the culture media. Culturing the leukocytes with LPS without n(CAT) significantly increases the production of TNF-a and IL-10 *(P* value 0.0001). Moreover, the cultures with n(CAT) show dramatically lower concentrations of TNF-a and IL-10 *(P* value 0.01 to 0.001), that are comparable with those of the control cells (resting leukocytes). This *ex vivo* study suggests that n(CAT) can downregulate the production of TNF-a and IL-10 by activated leukocytes, indicating a potential use of n(CAT) as an immunoregulator for hyperinflammation.

To further elucidate the immunoregulatory effect, leukocytes were cultured with injured HPAEpiC, of which cell injury was induced by H_2_O_2_ (Control #1, cell viability 85%). As shown in **Fig. 1J**, culturing the cells with leukocytes reduces the viability to 71%. Furthermore, adding 8, 16, and 40 μg/mL n(CAT) increases the viability to 82, 89, and 91%, respectively, which are comparable to those of Control #2 (leukocytes with untreated-HPAEpiC, 91% cell viability). This finding indicates that n(CAT) can not only protect, but also resuscitate, the injured alveolar cells, which is consistent with the observation presented in **Fig. 1G**. Furthermore, HPAEpiC was cultured with leukocytes activated by LPS. As shown in **Fig. 1K**, HPAEpiC (Blank) and HPAEpiC with LPS (Control #3) exhibit a similar cell viability, while HPAEpiC with LPS-activated leukocytes show a dramatically reduced cell viability of 67%. Moreover, adding 8, 16, and 40 μg/mL n(CAT) increases the cell viability to 78, 88, and 91%, respectively, which are comparable with those of Control #4 (un-activated leukocytes and HPAEpiC, cell viability 91%). This study suggests that n(CAT) can also protect healthy alveolar cells from injury by activated leukocytes, indicating an anti-inflammatory effect.

For therapeutic use, we first investigated the pharmacokinetics and biodistribution of n(CAT) in mice. For intravenous administration, BALB/c mice were administered 20 mg/kg of native catalase or n(CAT). **Fig. 2A** shows the biodistribution 6 h and 24 h post-injection; accumulation of n(CAT) is observed in the liver, kidney, lung, and lymph nodes, of which the average radiance is shown in **Fig. 2B**. **Fig. 2C** presents the pharmacokinetics, indicating that n(CAT) has a significantly longer circulation time than the native catalase. Based on the one-compartment model, n(CAT) exhibits a serum half-life of 8.9 h, which is 16.8-fold longer than the native CAT (0.5 h). Further analysis of the drug exposure time through the area under the curve (AUC) indicates that the mice that received n(CAT) had a significantly increased body exposure to catalase compared to the mice with native CAT (~ 2.5-fold increase) (**Fig. 2D**). The following were all within the normal ranges: the plasma levels of alanine aminotransferase, aspartate aminotransferase, and alkaline phosphatase (**fig. S3A**); the levels of urea and uric acid (**fig. S3B**); the total white blood cell (WBC) count; and the counts of lymphocytes, monocytes, and granulocytes (**fig. S3C**). Furthermore, H&E stained sections of the main organs do not show any noticeable tissue damage (**fig. S4**).

**Fig. 2.**
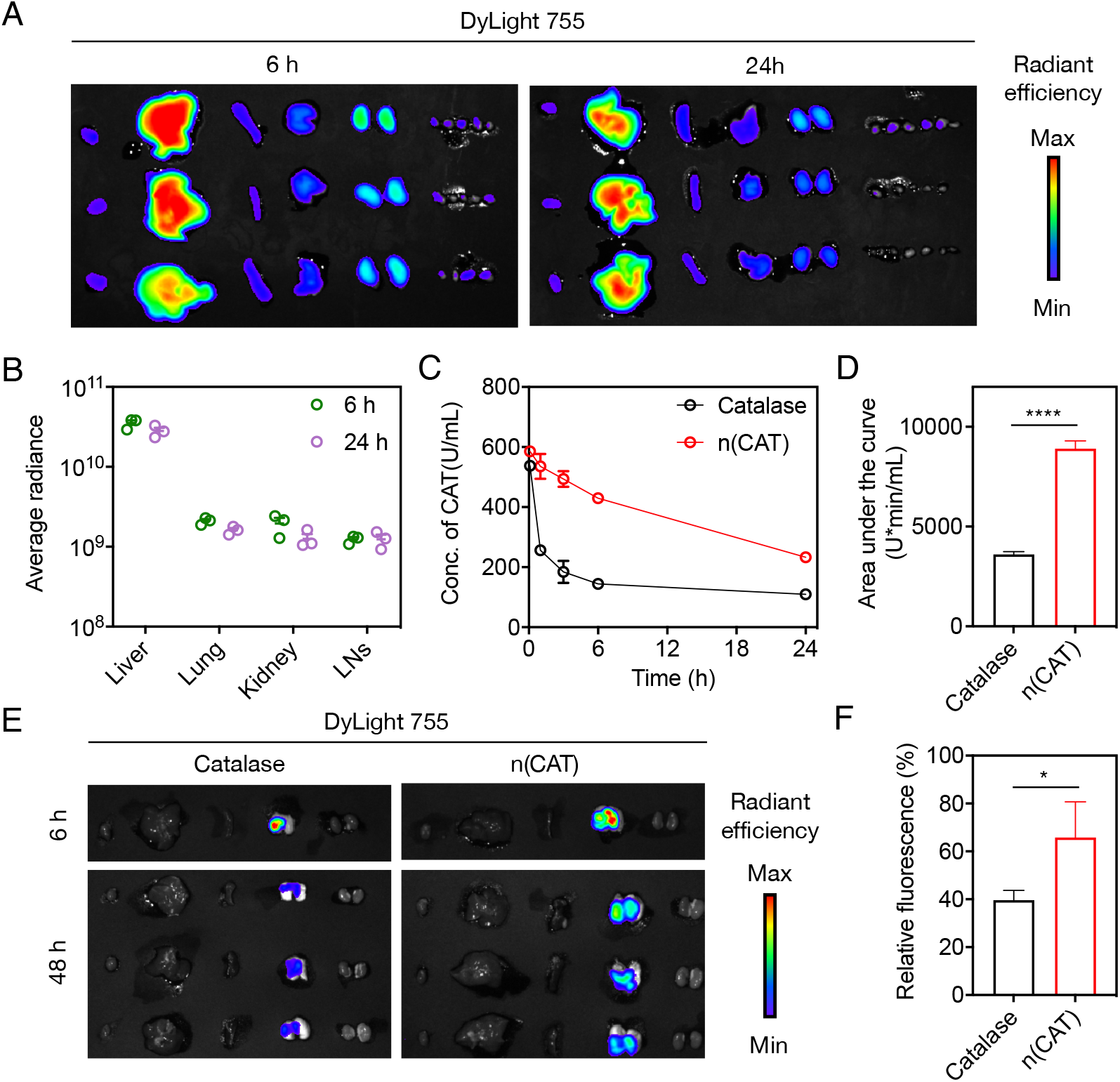
Pharmacokinetics and biodistribution of n(CAT) in mice. (A) Fluorescence imaging of the major organs, and (B) average radiance of n(CAT) in the liver, lung, and kidney 6 h and 24 h after intravenous administration of 20 mg/kg Cy7-labeled n(CAT). From left to right: heart, liver, spleen, lung, kidney, and lymph nodes. (C) Pharmacokinetics of native catalase and n(CAT) in BALB/c mice (n = 3) after intravenous administration of 20 mg/kg native catalase or n(CAT); blood samples were collected 0.1, 1, 3, 6, and 24 h after injection. (D) Drug exposure of the native catalase and n(CAT). (E) Fluorescence imaging of the major organs after intratracheal nebulization of native catalase and n(CAT). From left to right: heart, liver, spleen, lung, and kidney. (F) Relative fluorescence intensity of the lung 48 h after intratracheal nebulization of native catalase and n(CAT). *P* value: * < 0.05; **** < 0.0001.

For intratracheal nebulization, BALB/c mice were administered 2.5 mg/kg of native CAT or n(CAT) labeled with Alexa-Fluor-750. The mice receiving native catalase show fluorescent signal in the lung after 6 h, the intensity of which decreases significantly after 48 h. The mice receiving n(CAT) exhibit significantly higher fluorescent intensity after 6 h and 48 h **(Fig. 2E)**, which is confirmed by their fluorescent intensity plot after 48 h **(Fig. 2F)**. Except the lung, other organs (heart, liver, spleen, and kidney) after 48 h show negligible fluorescent signal, indicating that the as-administered n(CAT) was mainly retained within the lung. H&E stained sections of the main organs do not show any noticeable tissue damage (**fig. S5**).

The ability of n(CAT) to repress the replication of SARS-CoV-2 was examined in *rhesus macaques.* As illustrated in **Fig. 3A**, at day 0, all of the animals were inoculated with SARS-CoV-2 through the intranasal route. For the control group (C1, C2), two animals received 10 mL PBS though inhalation at day 2, 4, and 6, respectively. For the nebulization group, three animals (N1, N2, N3) received 5 mg of n(CAT) (10 mL) through inhalation at day 2, 4, and 6. For the intravenous group, two animals (I1, I2) received 10 mL PBS though inhalation and 5 mg/kg of n(CAT) intravenously at day 2, 4, and 6. Except N3 (sacrificed at day 21), the other animals were sacrificed at day 7.

**Fig. 3.**
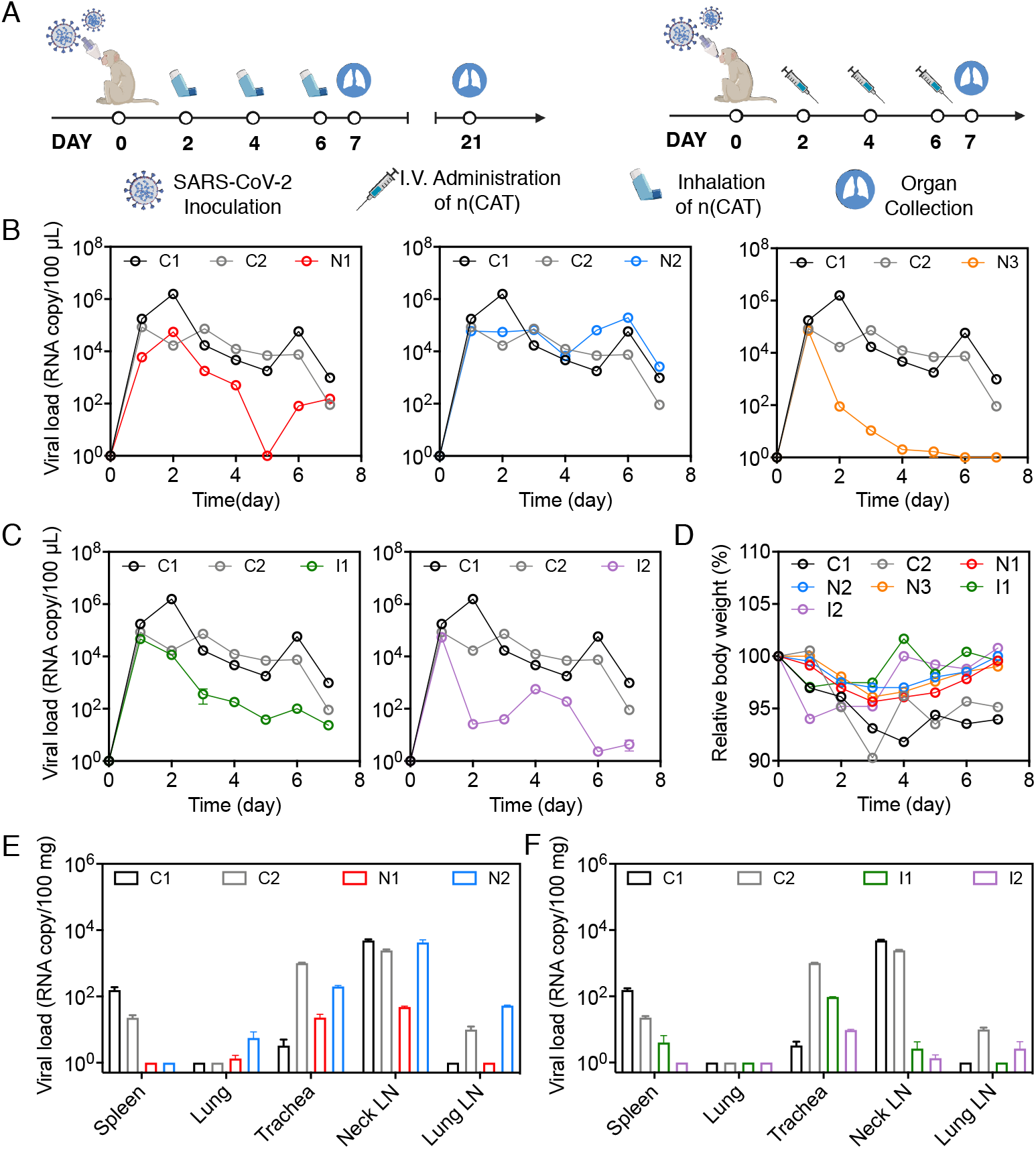
Ability of n(CAT) to repress the replication of SARS-CoV-2 in *rhesus macaques.* (A) Schematic showing the experiment design. (B, C) Viral loads in the nasal swabs of the animals that received (B) nebulization treatment (N1, N2, and N3) and (C) intravenous injection (I1 and I2) of n(CAT). (D) Relative bodyweight of the animals at day 1-7. (E, F) Viral loads in selective organs of the animals receiving (E) nebulization treatment and (F) intravenous injection of n(CAT) at day 7. Animals in the control group were marked as C1 and C2.

**Fig. 3B** shows the viral loads in nasal swabs for the control and nebulized group. N1 exhibits a viral load that is similar to C1 and C2 at day 1 and 2, after which the viral load rapidly decreases and becomes significantly lower than the control group. N3 shows a similar viral load to the control group at day 1, after which the viral load remains significantly lower than the control group. It is worth noting that the viral load of N3 at day 2 is lower than the control group. Nevertheless, the oral swabs confirmed that N3 was successfully infected, indicating an individual difference (**fig. S6**). N2 shows similar viral loads to the control group from day 1 to 7. **Fig. 3C** shows viral loads in the nasal swabs for the control and intravenous group. I1 exhibits a similar viral load to the control group at day 1 and 2, after which the viral load rapidly decreases and remains significantly lower than the control group. I2 also shows a similar viral load to the control group at day 1, after which the viral load remains significantly lower than the control group. Similarly, I2 shows a lower viral load than the control group, yet the oral swabs confirmed its active infection. **Fig. 3E-F** presents the viral RNA copy numbers in 100 mg of the organs, including lung, trachea, neck lymph node (LN), and lung LN. N1 shows significantly lower viral loads than the control group; whereas, N2 exhibits similar viral loads to the control group, which is consistent with the nasal-swab results. I1 and I2 show significantly lower viral loads than the control group, which is consistent with the nasal-swab results. No virus is detected from the organs of N3 (**fig. S7**). **Fig. 3D** shows the bodyweight change of the animals, suggesting that the experiment groups have less weight lost. These results confirm the ability of n(CAT) to repress the replication of SARS-CoV-2 in *rhesus macaques.*

**Fig. 4A-F** shows the liver and renal functions of the control and experimental group, which exhibit similar levels of alanine aminotransferase, aminotransferase aspartate aminotransferase, alkaline phosphatase, albumin, uric acid, creatine, and blood urea nitrogen, indicating that intravenous administration of 5 mg/kg of n(CAT) did not cause any noticeable liver or renal toxicity. Meanwhile, all of the groups show similar blood routine and other indexes for liver function (**fig. S8**). Similar results were also observed in healthy *rhesus macaques* inhaling 2.0 mg/kg (**fig. S9)** n(CAT) per day for 7 d, suggesting that n(CAT) does not cause noticeable liver or kidney toxicity.

**Fig. 4.**
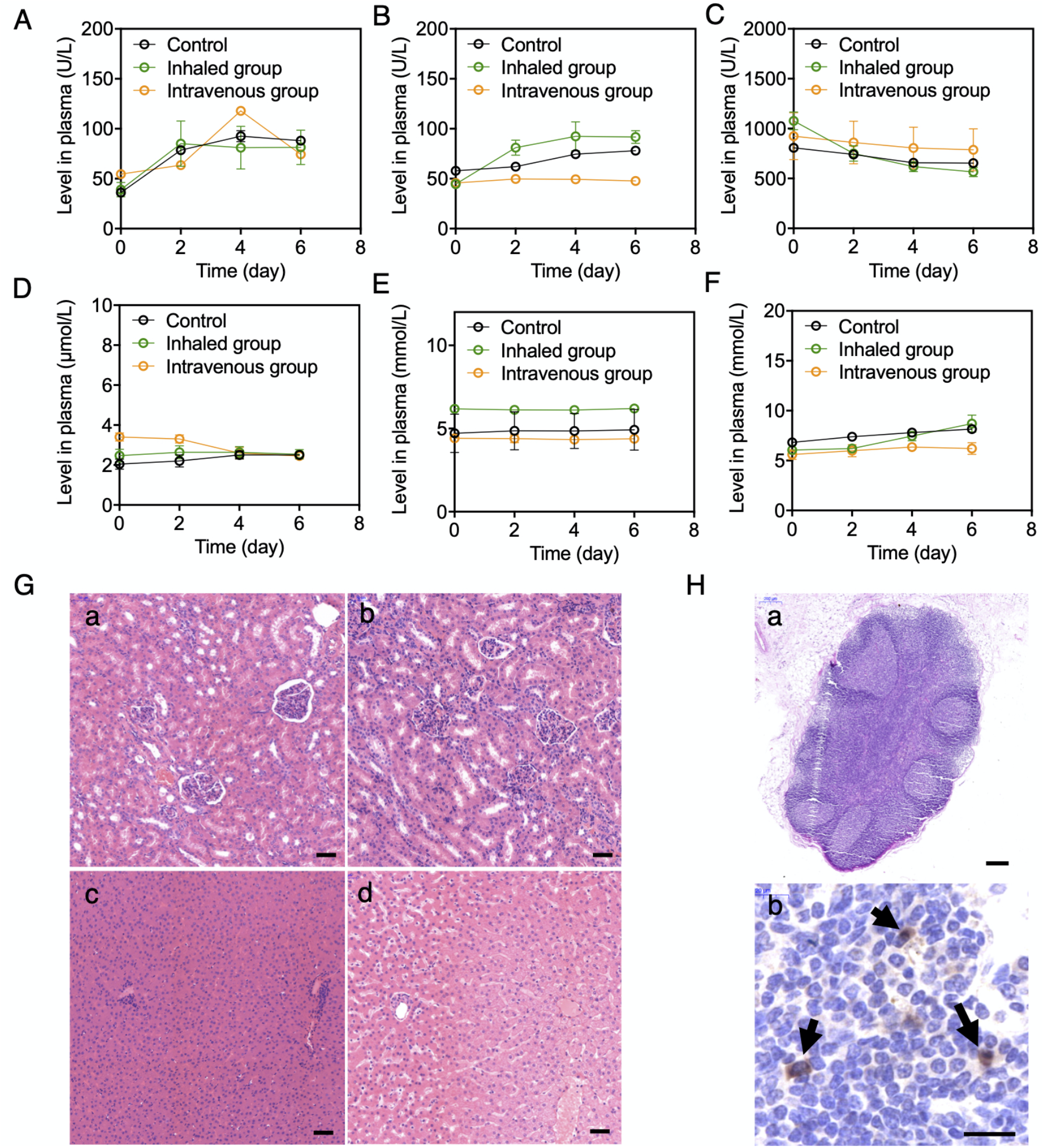
Biosafety and histology of SARS-CoV-2 infected *rhesus macaques.* (A) Aspartate aminotransferase (AST), (B) alanine aminotransferase (ALT), (C) alkaline phosphatase (ALP), (D) uric acid (UA), (E) urea, and (F) blood urea nitrogen (BUN) levels of the animals in the control, inhaled, and intravenous groups. (G) H&E stained sections of the kidneys (a, b) and livers (c, d) in the control (a, c) and inhaled groups (b, d) (scale bar = 50 μm). (H) (a) Representative H&E stained section (scale bar = 200 μm) and (b) immunohistochemistry staining of SARS-CoV-2 nucleocapsid protein [6H3], demonstrating scattered positive mononuclear cells (arrows) within the lung LN in C1 (scale bar = 20 μm).

**Fig. 4G** presents representative H&E sections of kidneys (a, b) and liver (c, d) from animals in the control (a, c) and inhaled group (b, d). The kidneys show neither evidence of interstitial nephritis nor acute tubular injury; the livers exhibit neither steatosis, hepatocyte necrosis, inflammation, cholestasis, nor bile duct injury. Histologic sections of lung tissues in both the control and inhaled groups exhibit unremarkable alveolar architecture, with no evidence of acute lung injury in the form of hyaline membranes, intra-alveolar fibrin, organizing pneumonia, or reactive pneumocyte hyperplasia. The airway epithelium is unremarkable. Vascular compartments are free of thrombi **(fig. S10)**. There is no evidence of eosinophilia or vasculitis, and no viral cytopathic effect is identified. The H&E staining of other major organs also shows no tissue injury for both the control and inhaled group (**fig. S11)**, confirming the biosafety of n(CAT) administered through intravenous injection or inhalation. In addition, **Fig. 4H** also presents a representative H&E section (a) and immunohistochemistry for SARS-CoV-2 nucleocapsid protein (b) of the lung LN in one animal from the control group (C1). Reactive follicular hyperplasia could be observed in the H&E section, and scattered positive mononuclear cells (black arrows) indicate the SARS-CoV-2 infection in the lymph node.

The action mechanism of n(CAT) is unclear. In addition to being a weapon against pathogens, ROS also serve as signaling molecules in numerous physiological processes.(*33*) For example, it has been documented that H_2_O_2_ generation after wounding is required for the recruitment of leukocytes to the wound(*34*), and ROS is necessary for the release of pro-inflammatory cytokines to modulate an appropriate immune response(*22*). Eliminating the H_2_O_2_ excessively produced during inflammation also minimizes the downstream ROS, which assists to downregulate production of cytokines, mitigate recruitment of excessive leukocytes, and repress replication of the viruses. It is also worth noting that immunosuppressive steroids, such as prednisone and dexamethasone, are proven to be effective for treatment of hyperinflammation in severe COVID-19 patients(*9*). Glucocorticoids constitute powerful, broad-spectrum antiinflammatory agents that regulate cytokine production, but their utilization is complicated by an equally broad range of adverse effects(*35, 36*). For instance, in a retrospective study of 539 patients with SARS who received corticosteroid treatment, one-fourth of the patients developed osteonecrosis of the femoral head(*37*). We speculate that n(CAT) could also regulate cytokine production, but through a different pathway – reinstating immune homeostasis through eliminating excessively produced ROS.

In conclusion, we have shown the anti-inflammatory effect and ability of catalase to regulate cytokine production in leukocytes, protect alveolar cells from oxidative injury, and repress the replication of SARS-CoV-2 in *rhesus macaques* without noticeable toxicity. Moreover, it is worth noting that catalase is safe and commonly used as a food additive and dietary supplement, and that pilot-scale manufacturing of n(CAT) has been successfully demonstrated. In contrast to the current focus on vaccines and antiviral drugs, this may provide an effective therapeutic solution for the pandemic, as well as treatment of hyperinflammation in general.

