## Supplementary Information for "An Antioxidant Enzyme Therapeutic for COVID-19"

**This PDF file includes:**

Materials and methods

Fig. S1 to S11

References for supplementary information

### Materials and Methods

#### 1. Materials

Catalase, trypsin, N-acryloxysuccinimide (NAS), 2-methacryloyloxyethyl phosphorylcholine (MPC), N-(3-Aminopropyl)methacrylamide hydrochloride (APM), Bis-acrylamide (BIS), N,N,N′,N′-tetramethylethylenediamine (TEMED), and ammonium persulfate (APS) were purchased from Sigma Aldrich. Sulfo-Cy7-NHS was purchased from Beijing Fanbo Biochemicals Co., Ltd. Lipopolysaccharides (LPS) was purchased from Solarbio, Beijing. Alexa Fluor 750 NHS ester (Alexa Fluor-750-NHS) was purchased from Thermo Fisher Scientific. Human pulmonary alveolar epithelial cells (HPAEpiC) were purchased from Bnbio.

#### 2. Synthesis and characterization of n(CAT)

Catalase was dissolved in phosphate buffered saline (PBS), dialyzed overnight against PBS at 4 ˚C, and filtered using a 0.22 $\mu$m filter. As-purified catalase was first conjugated with acrylate moieties by reacting with NAS (20:1, n/n, NAS:Catalase) at 4 ˚C for 2 h, and dialyzed overnight against PBS at 4 ˚C. To synthesize n(CAT), MPC, APM, and BIS (21600:2400:2400:1, n/n, MPC:APM:BIS:Catalase) were added to form a solution, and then APS (2400:1, n/n, APS:Catalase) and TEMED (2:1, w/w, TEMED:APS) were added to initiate the polymerization. After reacting at 4 ˚C for 2 h, the solution was dialyzed against PBS and passed through a phenyl-sepharose CL-4B column (GE) using 10x PBS as the elusion buffer. The concentration of n(CAT) was determined with BCA protein assay. The size distribution and zeta potential were measured by dynamic light scattering (DLS, Zetasizer Nano Instruments). The diameter and morphology were evaluated by TEM (Tecnai T12).

#### 3. Quantification of enzyme activity and stability

Native catalase or n(CAT) was added to solutions containing 1.25, 2.50, 5.00, and 10.00 mM $H_2O_2$ at a final catalase concentration of 1 $\mu$g/mL, respectively. The reaction rates were measured by the absorption at 240 nm using UV-VIS (Beckman) every 10 s (40 s in total). To determine the thermal stability, native catalase or n(CAT) was diluted with PBS to a final concentration of 0.1 mg/mL, and incubated at 37 °C for 24 h. The activities of the enzymes were measured by adding the incubated catalase or n(CAT) to 10 mM $H_2O_2$ at a final catalase concentration of 1 $\mu$g/mL. To measure the proteolytic stability, native catalase or n(CAT) was diluted with PBS to a final concentration of 0.1 mg/mL, after which trypsin (Sigma Aldrich) was added to the solutions at a final concentration of 50 $\mu$g/mL, and incubated at 37 °C for 2 h. The enzyme activities were tested in 1000 $\mu$M $H_2O_2$ at a final catalase concentration of 1 $\mu$g/mL.

#### 4. *In vitro* and *ex vivo* studies

Human pulmonary alveolar epithelial cells (HPAEpiC) were cultured on 25 cm$^2$ tissue culture flasks containing RPMI1640 media, supplemented with 10% FBS and 1% P/S. HPAEpiC were seeded in 96 well-plates at 5,000 cells/well 12 h prior to *in vitro* tests, and were seeded at 10,000 cells/well 12 h before the *ex vivo* studies. The cell viability was tested by the CCK-8 Kit (Dojindo) following the protocol provided.

The cytotoxicity of n(CAT) was examined by culturing HPAEpiC with 20, 10, 200, and 1000 $\mu$g/mL of n(CAT) for 12 h. To validate the resuscitative ability of n(CAT) for injured cells, $H_2O_2$ was added to HPAEpiC at a final concentration of 1000 $\mu$M, and incubated for 24 h. The media was then removed, and the cells were washed with chilled PBS for three times. n(CAT) was then added at final concentration of 2, 4, 8, 16, and 20 $\mu$g/mL, respectively, and incubated for another 12 h. To validate the ability of n(CAT) to prevent oxidative injury, HPAEpiC were pre-incubated with 2, 4, 8, 16, and 20 $\mu$g/mL of n(CAT) for 12 h, respectively, after which $H_2O_2$ was added to the cells at a final concentration of 1000 $\mu$M, and incubated for 24 h. The cell viability tests were conducted after all of the *in vitro* tests.

Leukocytes were separated from whole blood from a donor following a protocol provided by Thermo Fisher Scientific. To confirm the ability of n(CAT) to regulate cytokine production, leukocytes were seeded in a 96 well-plate at 100,000 cells/well, and LPS was added at a final concentration of 1 $\mu$g/mL to activate the leukocytes. n(CAT) was added at a final concentration of 2, 4, 8, 16, and 20 $\mu$g/mL, respectively, and incubated for 12 h. The concentration of TNF-α and IL-10 in the media was measured with an enzyme-linked immunosorbent assay kit (Abcam) following the protocols provided.

To validate the ability of n(CAT) to protect injured cells against leukocytes, $H_2O_2$ was added to HPAEpiC at a final concentration of 500 $\mu$M, and incubated for 12 h to mimic cell injury. The injured cells were then washed three times with chilled PBS, followed by adding fresh leukocyte-containing media (100,000 cells in each well). n(CAT) was added at a final concentration of 2, 4, 8, 16, and 20 $\mu$g/mL, respectively, and incubated for another 12 h. PBS was added to the injured HPAEpiC as Control #1, and leukocytes were added to HPAEpiC without $H_2O_2$ treatment as Control #2.

To confirm the ability of n(CAT) to protect HPAEpiC against injury from activated leukocytes, HPAEpiC were incubated with 100,000 leukocytes containing 1 $\mu$g/mL lipopolysaccharides (LPS) to activate the leukocytes. n(CAT) was then added at final concentration of 2, 4, 8, 16, and 20 $\mu$g/mL, respectively, and incubated for 24 h prior to measuring the cell viability. 1 $\mu$g/mL LPS was added to HPAEpiC as Control #3, and leukocytes were added to HPAEpiC without LPS as Control #4.

**5. Biodistribution and pharmacokinetics in mice**

To assess the biodistribution of n(CAT), native catalase and n(CAT) labeled with sulfo-Cy7-NHS or Alexa Fluor-750-NHS were used for intravenous injection and intratracheal instillation, respectively. To achieve the labeling, sulfo-Cy7-NHS was added to native catalase or n(CAT) (5:1, n/n, Cy7:CAT), and the reaction was kept at 4 °C overnight. The unreacted Cy7 was removed by dialysis against PBS overnight at 4 °C, and the concentrations of catalase and n(CAT) were determined by BCA assay, respectively. To label native catalase and n(CAT) with Alexa Fluor-750-NHS, a similar protocol was used.

BALB/c mice (8 w, 22±2 g, n=4) received 2.5 mg/kg of labeled native catalase or n(CAT) through intratracheal instillation, and were euthanized 6 h and 48 h post-instillation. BALB/c mice (8 w, 22±2 g, n=3) received 20 mg/kg of labeled native catalase or n(CAT) through tail-vein injection, and were euthanized 6 h and 24 h post-injection. The organs of these animals (heart, liver, spleen, lung, and kidney) were collected and imaged with an *in vivo* imaging system (IVIS spectrum, Perkin Elmer). The organs were also fixed in 10% neutral-buffered formalin and embedded in paraffin. The sections (4 $\mu$m in thickness) were stained with haematoxylin and eosin, and examined with a light-microscope.

To evaluate the pharmacokinetics, BALB/c mice (8 w, 22±2 g, n=3) received 20 mg/kg native catalase or n(CAT) through tail-vein injection, and blood samples were collected at 0.1, 1, 3, 6, and 24 h post-injection. The serum was separated from the whole blood by centrifugation at 4000 x g for 10 min, and the serum catalase activity was assessed by a microplate-based method according to a published test with a minor modification(*1*). Briefly, the serum was diluted five times with 50 mM phosphate buffer (PB, pH 7.4). Subsequently, 20 $\mu$L of the diluted serum was added to 100 $\mu$L $H_2O_2$ (50 mM) and incubated for 60 s. The reaction was terminated by adding 100 $\mu$L ammonium molybdate solution (50 mM), and the absorbance at 405 nm was measured by a UV-VIS (Beckman) and recorded as $A_{test}$. Three blank samples were also tested as controls. First, 100 $\mu$L $H_2O_2$ (50 mM) was mixed with 100 $\mu$L ammonium molybdate solution (50 mM), followed by adding 20 $\mu$L of the diluted serum. The absorbance at 405 nm was recorded as $A_1$. Second, 120 $\mu$L PB (50 mM) was mixed with 100 $\mu$L ammonium molybdate solution (50 mM), and the absorbance was recorded as $A_2$. Finally, 100 $\mu$L $H_2O_2$ (50 mM) was mixed with 20 $\mu$L PB (50 mM) and 100 $\mu$L ammonium molybdate solution (50 mM), and the absorbance was recorded as $A_3$. The activity of catalase in the serum was then calculated by the following equation: catalase activity (U/mL) = $\frac{A_{test}-A_1}{A_3-A_2} \times 5$.

### 6. Studies in SARS-CoV-2 infected *rhesus macaques*

The studies were conducted in a BSL-3 laboratory in the Institute of Medical Biology, Chinese Academy of Medical Sciences (BSL number SWAQ20200404, animal protocol number DWSP202001020). SARS-CoV-2 (SARS-Cov-2/KM 1/2010) was originally separated from the sputum of a patient infected with COVID-19 in Kunming, Yunnan, China. Seven *rhesus macaques* (female, 10-12 mo., body weight 1.5-2.5 kg) were randomly distributed into three groups: control (n=2, C1, C2), inhaled (n=3, N1, N2, N3), and intravenous (n=2, I1, I2).

All of the animals were inoculated with SARS-CoV-2 under anesthesia through intranasal route (100 $\mu$L per nostril) with a suspension of $10^5$ CCID50 SARS-CoV-2 in PBS. The animals in the control group inhaled 10 mL PBS at day 2, 4, and 6 post-inoculation (p.i.) with a nebulizer (OMRON NB-150U). The animals in the inhaled group received 5 mg n(CAT) (10 mL) at day 2, 4, and 6 p.i. with the nebulizer. The animals in the intravenous group received 5 mg/kg n(CAT) at day 2, 4, and 6 p.i. through saphenous vein injection. The animals in the intravenous treatment group also received 10 mL PBS at day 2, 4, and 6 p.i. with the nebulizer. All of the animals were anesthetized daily with ketamine (5 mg/kg) during day 1-7 p.i. Nasal swabs and oral swabs were collected with cotton swabs from the nasal and oral cavity, respectively. Body weights were also measured daily. For each animal, 1 mL blood sample was collected from the saphenous vein with anti-coagulation tubes, and another 2 mL blood sample was collected with pro-coagulation tubes. The serum was then separated for biochemical tests and blood routine tests with a Mindray BS-200 chemistry analyzer and a Sysmex XT 2000i automated hematology analyzer, respectively. Except N3 (sacrificed at day 21 p.i.), all animals were sacrificed at day 7 p.i. Autopsies of the animals were performed according to a standard protocol, and the organs were collected.

#### 6.1 Viral load testing

To extract the viral RNA samples from nasal and oral swabs, the swabs were added to 1 mL TRNzol and mixed by vortex; 200 $\mu$L of the liquid was then mixed with 600 $\mu$L TRNzol. To extract the viral RNA samples from tissues, as-collected tissues were homogenized (SCIENTZ-48 homogenizer) with TRNzol (tissue: 20 wt-%). After homogenization, 200 $\mu$L homogenate was mixed with 600 $\mu$L TRNzol. Chloroform (200 $\mu$L) was added to each of the mixtures, placed at room temperature for 10 min, and centrifuged at 12,000 rpm for 10 min. After centrifugation, 500 $\mu$L supernatant was transferred to a tube. Isopropanol (500 $\mu$L) was then added, placed at room temperature for 10 min, and centrifugated at 12,000 rpm for 10 min. The resultant precipitation was then mixed with ethanol (1 mL, 75 vol-%), and centrifugated at 10,000 rpm for 5 min at 4 °C. After removal of the ethanol, the precipitation was placed at room temperature for 5 min and dissolved with 30 $\mu$L RNase-free DI water. The SARS-CoV-2 RT-qPCR was performed following a previously published protocol(*2*).

**6.2 Histology and immunohistochemistry staining**

Tissues for light-microscope examination were fixed in 10% neutral-buffered formalin, embedded in paraffin, and 4 $\mu$m sections were stained with haematoxylin and eosin. In the immunohistochemistry study, paraffin was removed from the sections, and the sections were treated with citric acid buffer (pH 6.0) for 20 min at 100 °C. Endogenous peroxidase was then blocked with 3% hydrogen peroxide for 25 min at room temperature. The sections were briefly washed with PBS (pH 7.4) for three times, and were blocked with 3% bovine serum albumin (BSA) for 30 min at room temperature. The slides were then incubated with a rabbit polyclonal antibody against SARS-CoV diluted 1:100 with PBS, and kept at 4 °C overnight. After washing with PBS for three times, the slides were incubated with horseradish peroxidase (HRP) labeled goat-anti-rabbit IgG diluted 1:200 with PBS for 50 min at room temperature. After washing with PBS, HRP activity was revealed by adding freshly prepared DAB solution, and the reaction was terminated by washing with DI water when a positive brown color was observed under the light-microscope. The sections were counterstained with haematoxylin.

**7. Statistical Analysis**

All results are presented as the mean ± standard error of the mean (s.e.m.) as indicated. Paired t-tests and one-way ANOVA were used for multiple comparisons (when more than two groups were compared). All statistical analyses were conducted with Prism Software (Prism 8.0).

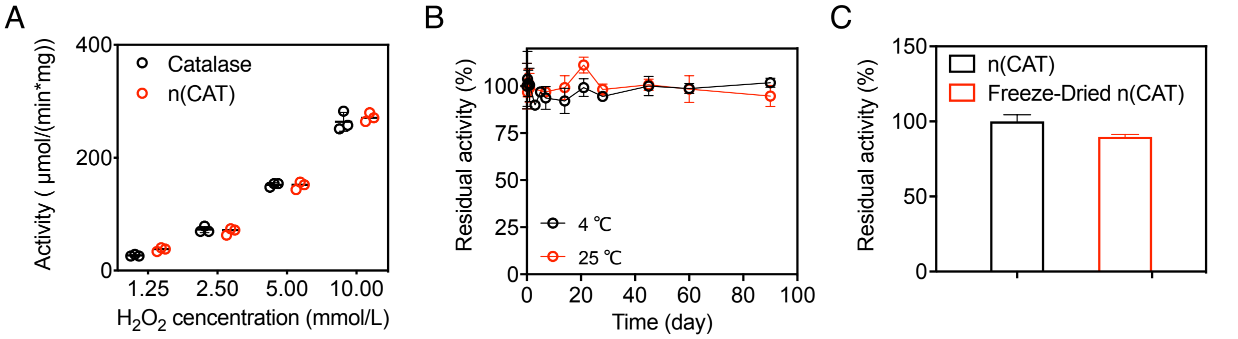


**Fig. S1.** The activity and stability of native catalase and n(CAT). (A) Enzyme activity of n(CAT) in different concentrations of $H_2O_2$. (B) Residual activity of n(CAT) in solution stored at 4 °C and 25 °C for 90 days. (C) Residual activity of n(CAT) after freeze-drying.

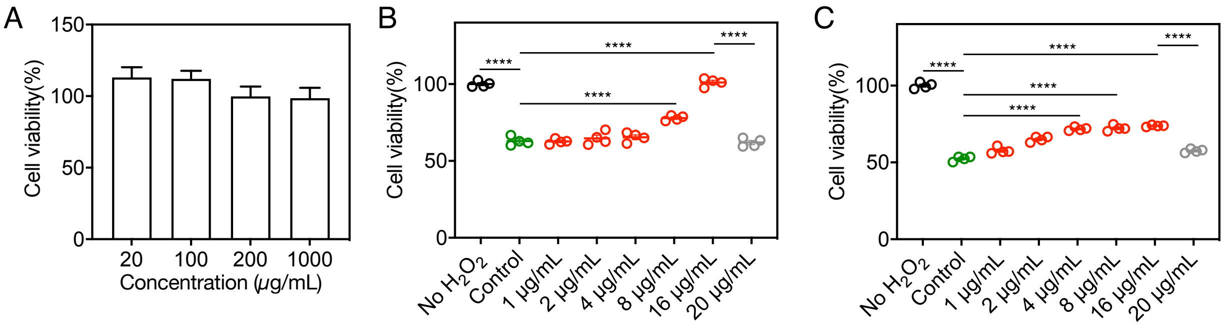


**Fig. S2.** Cytotoxicity, protective and resuscitative effects of n(CAT) in HPAEpiC against oxidative injury. (A) Cell viability of HPAEpiC in the presence of different concentrations of n(CAT) for 12 h. (B) Cell viability of HPAEpiC pre-cultured with different concentrations of n(CAT) for 12 h followed by addition of $H_2O_2$ (1000 mM) and culturing for 24 h. 20 $\mu$g/mL native catalase was used as a control. (C) Cell viability of HPAEpiC pre-cultured with 1000 mM $H_2O_2$ for 24 h, followed by culturing in a fresh media containing different concentrations of n(CAT) for 12 h. 20 $\mu$g/mL native catalase was used as a control. P value: **** < 0.0001.

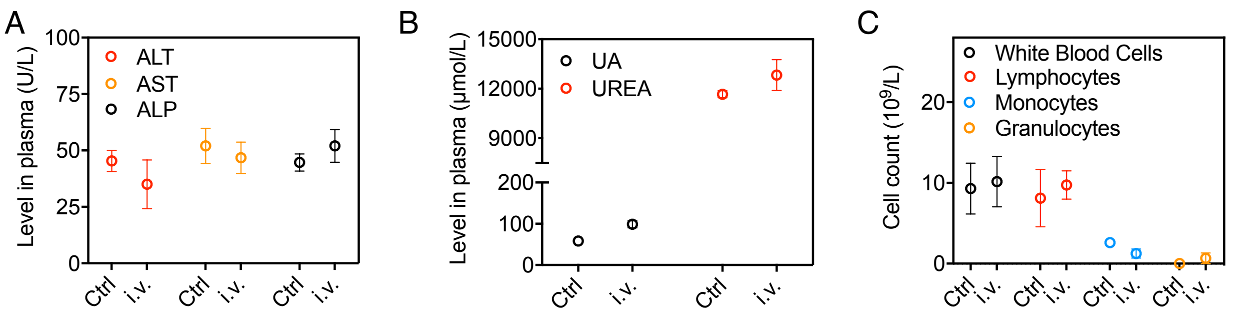


**Fig. S3.** Evaluation of the biosafety of n(CAT) in BALB/c mice. (A) Plasma levels of alanine aminotransferase (ALT), aspartate aminotransferase (AST), and alkaline phosphatase (ALP), (B) urea (UREA) and uric acid (UA), and (C) blood routine 24 h after intravenous injection of 20 mg/kg n(CAT).

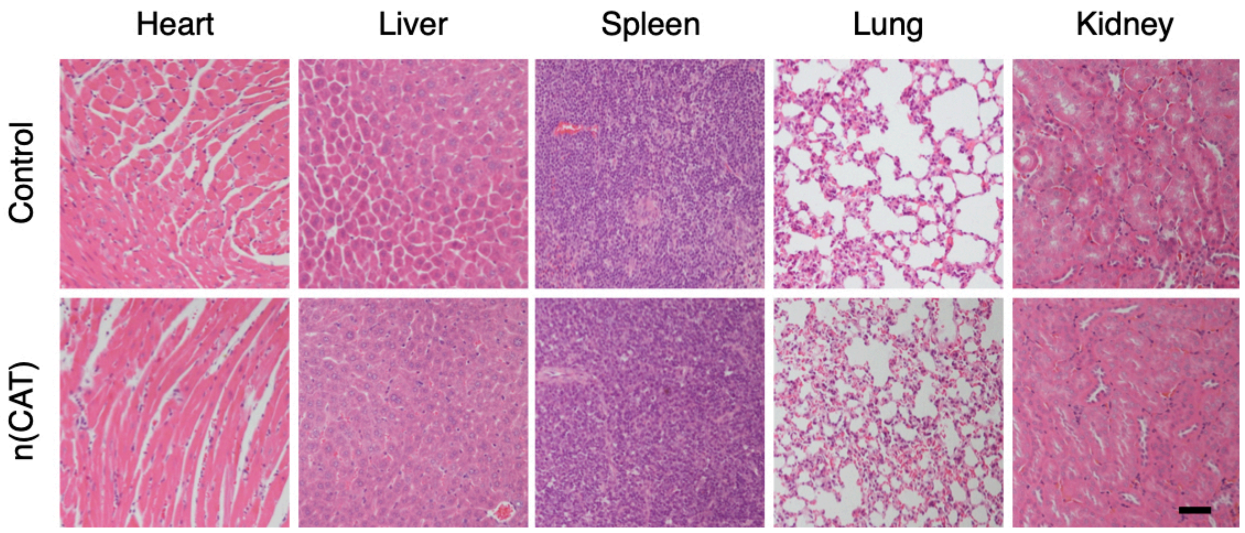


**Fig. S4**. Representative H&E staining sections of major organs in BALB/c mice 24 h after intravenous injection of 20 mg/kg n(CAT). (scale bar = 50 $\mu$m)

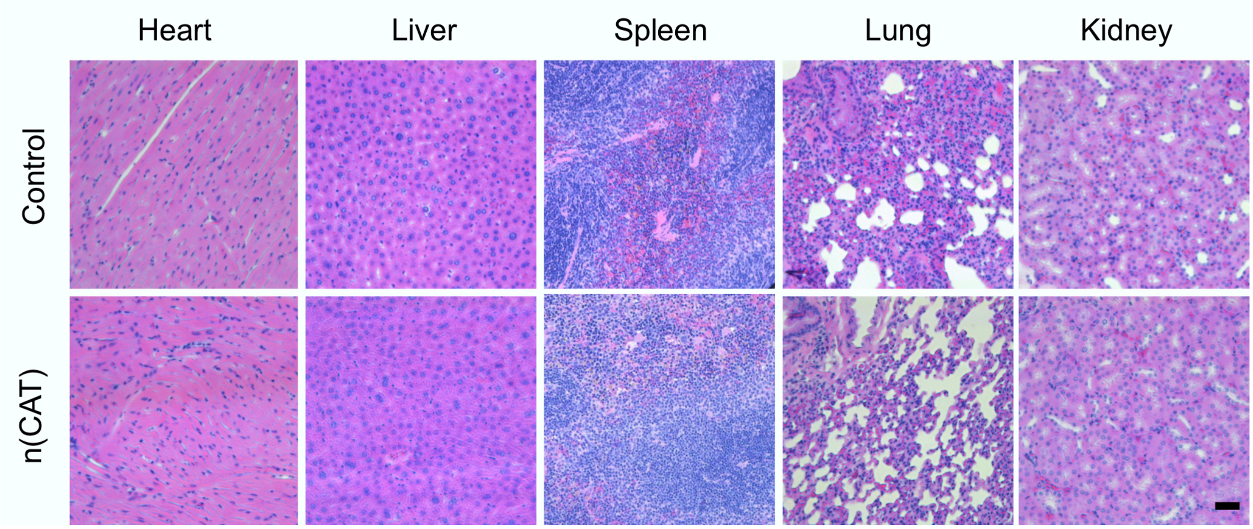


**Fig. S5.** Representative H&E staining sections of major organs in BALB/c mice 24 h after intratracheal instillation of 2.5 mg/kg n(CAT). (scale bar = 50 $\mu$m)

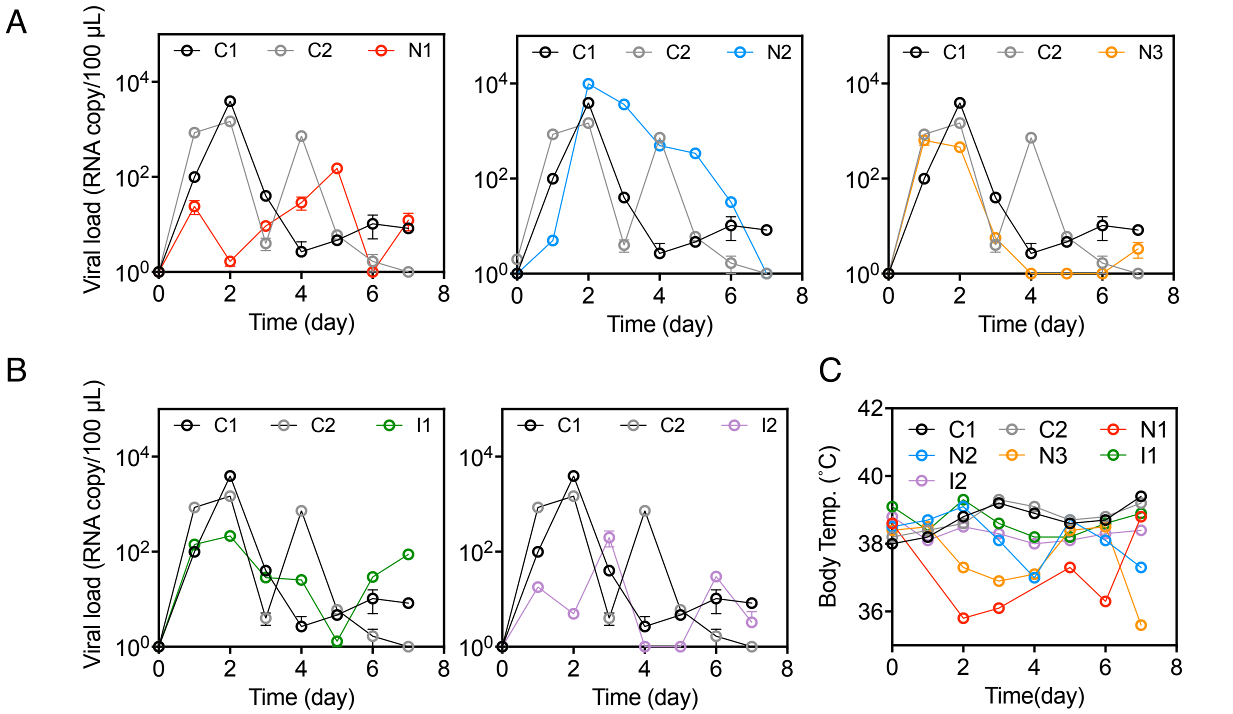


**Fig. S6.** Viral loads in oral swabs and body temperature change in SARS-CoV-2 infected rhesus macaques. (A) Viral loads in the orals swabs in the inhaled group. (B) Viral loads in the oral swabs in the intravenous group. (C) Body temperature change in all the animals.

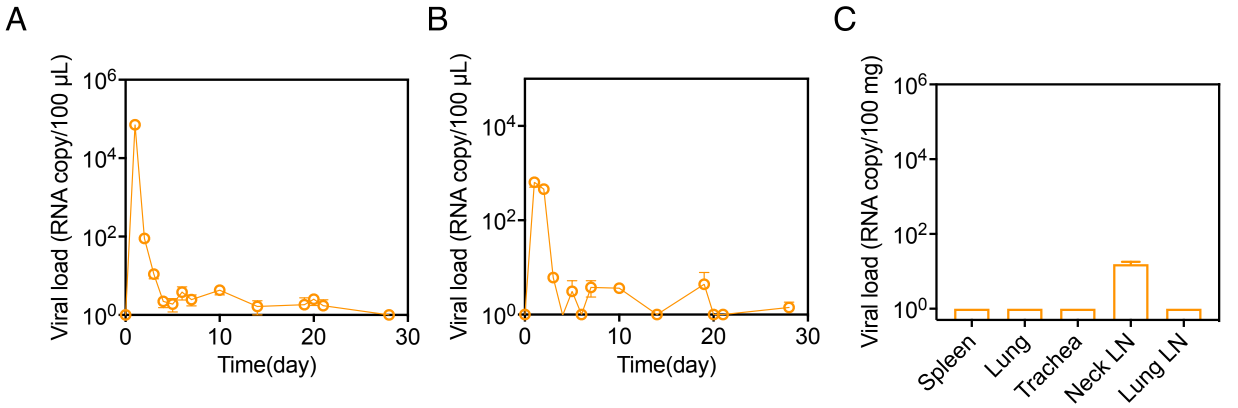


**Fig. S7.** Viral loads of N3 in the inhaled group at day 1-28 p.i. (A) Viral loads in the nasal swabs. (B) Viral loads in the oral swab. (C) Viral loads in major organs at day 28 p.i.

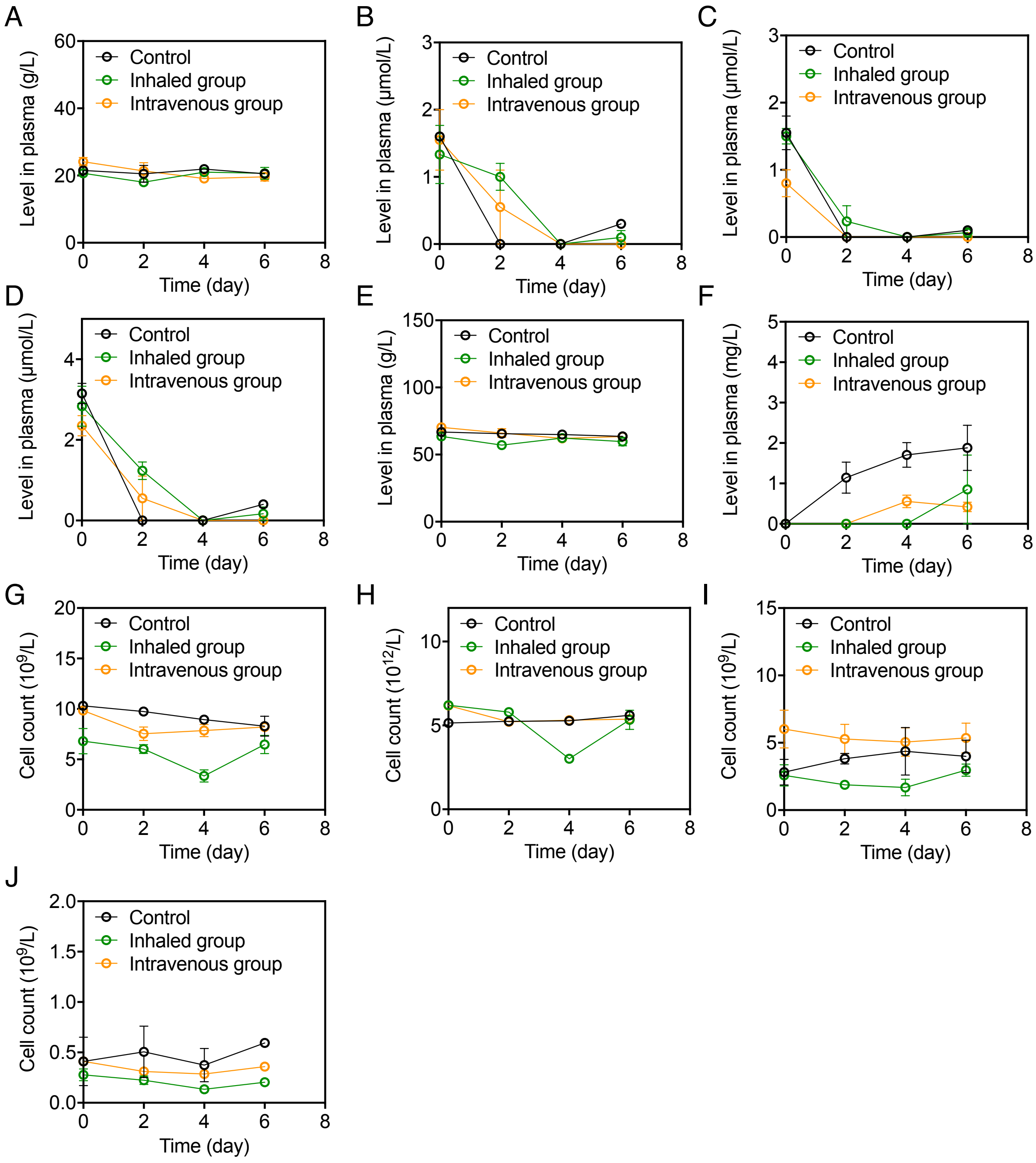


**Fig. S8.** Evaluation of the biosafety of n(CAT) in SARS-CoV-2 infected rhesus macaques. (A) Globulin (GLB); (B) indirect bilirubin (IBIL); (C) direct bilirubin (DBIL); (D) total bilirubin (TBIL); (E) total protein (TP); (F) C-reactive protein (CRP); (G) white blood cell (WBC) number; (H) red blood cell (RBC) number; (I) neutrophil number; and (J) monocyte number tests at day 1-7 p.i. in the control, inhaled, and intravenous groups.

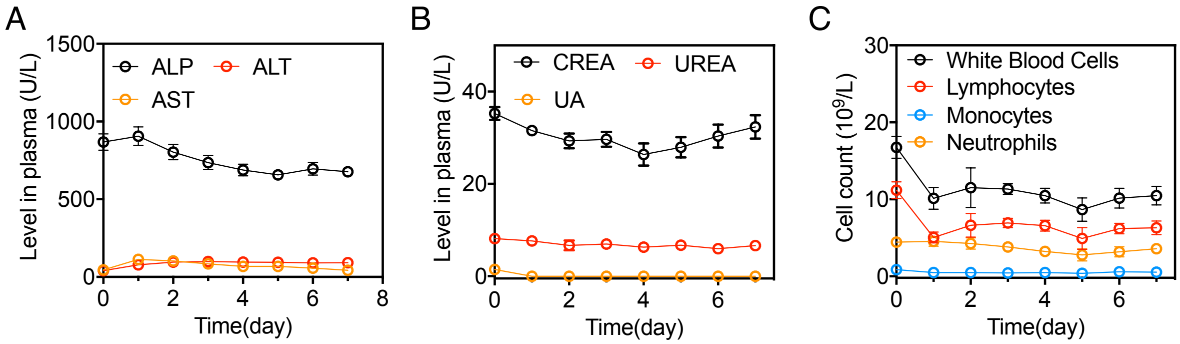


**Fig. S9.** Evaluation of the biosafety in healthy rhesus macaques receiving 2 mg/kg n(CAT) through inhalation daily for seven days. (A) Liver (AST, ALT, ALP) functions, (B) renal functions (UA, urea. creatine (CREA)), and (C) blood routine examination.

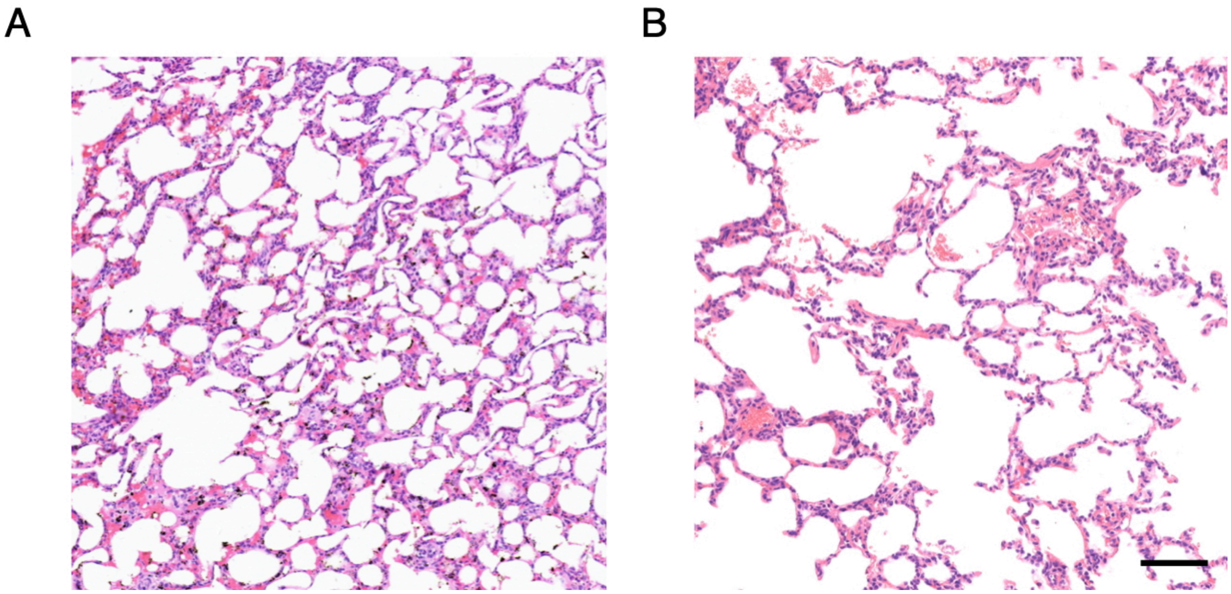


**Fig. S10.** Representative H&E staining sections of lungs in the (A) control group and (B) inhaled group. (scale bar = 100 $\mu$m)

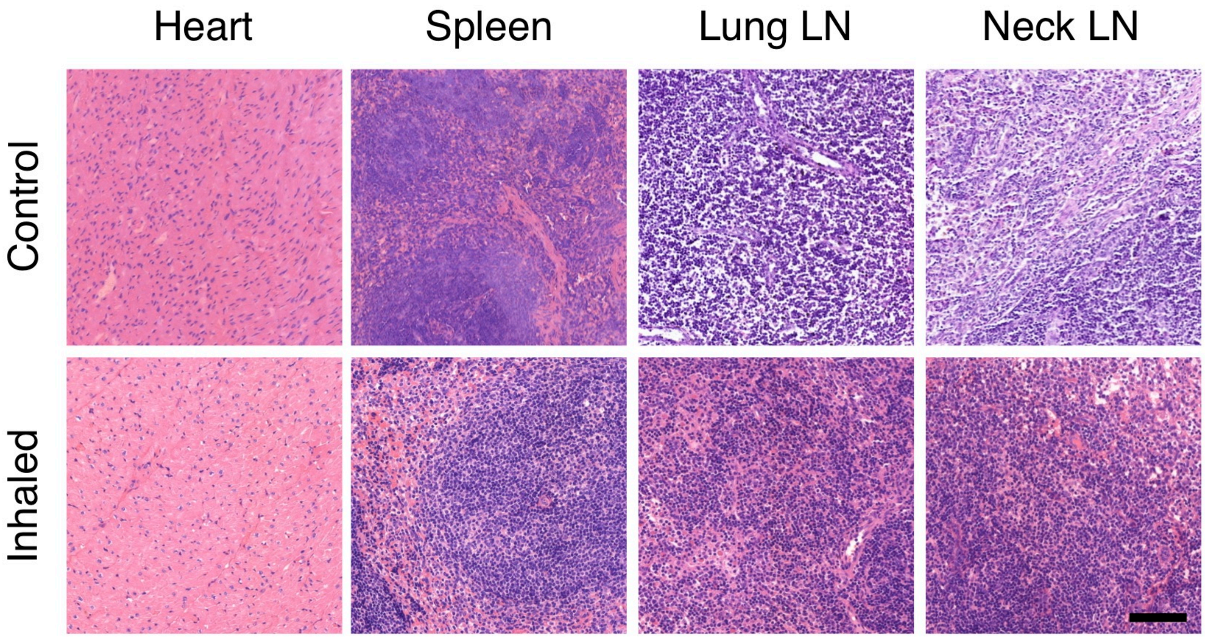


**Fig. S11.** Representative H&E staining sections of other major organs in the control and inhaled groups. (scale bar = 100 $\mu$m)
